# Individual differences in neural event segmentation of continuous experiences

**DOI:** 10.1101/2022.09.09.507003

**Authors:** Clara Sava-Segal, Chandler Richards, Megan Leung, Emily S. Finn

**Affiliations:** Department of Psychological and Brain Sciences Dartmouth College Hanover, NH 03748, USA; Section on Functional Imaging Methods, Laboratory of Brain and Cognition National Institute of Mental Health, Bethesda, MD, 20892-9663, USA

**Keywords:** event segmentation, fMRI, individual differences, naturalistic stimuli, stimulus properties

## Abstract

Event segmentation is a spontaneous part of perception, important for processing continuous information and organizing it into memory. While neural and behavioral event segmentation show a degree of inter-subject consistency, meaningful individual variability exists atop these shared patterns. Here we characterized individual differences in the location of neural event boundaries across four short movies that evoked variable interpretations. Event boundary alignment across subjects followed a posterior-to-anterior gradient that was tightly correlated with the rate of segmentation: slower-segmenting regions that integrate information over longer time periods showed more individual variability in boundary locations. This relationship held irrespective of the stimulus, but the degree to which boundaries in particular regions were shared versus idiosyncratic depended on certain aspects of movie content. Furthermore, this variability was behaviorally significant in that similarity of neural boundary locations during movie-watching predicted similarity in how the movie was ultimately remembered and appraised. In particular, we identified a subset of regions in which neural boundary locations are both aligned with behavioral boundaries during encoding and predictive of stimulus interpretation, suggesting that event segmentation may be a mechanism by which narratives generate variable memories and appraisals of stimuli.

## Introduction

Event segmentation refers to the spontaneous chunking of continuous experiences into meaningful distinct units or *events* as part of everyday perception (Zacks and Swallow 2007; Zacks et al. 2007). This cognitive mechanism is an automatic and adaptive part of perceptual processing, optimizing attention, and facilitating the subsequent organization of experiences into memory (Zacks and Swallow 2007; Zacks et al. 2007; Kurby and Zacks 2008; DuBrow and Davachi 2016; McGatlin et al. 2018).

Event boundaries, typically reported behaviorally by pressing a button to indicate transitions between events (Newtson 1973), are to some extent “normative”, or consistent across people. Boundary locations not only have high intersubject reliability, but are also meaningfully structured, corresponding to moments of decreased contextual stability where the prediction of the immediate future fails, such as changes in action, goals, or locations (Newtson et al. 1976; Speer et al. 2009; Zacks et al. 2009; Raccah et al. 2022). Neural responses at normative event boundaries have been reported in several regions including the hippocampus, lateral frontal cortex, and medial and lateral posterior cortex (Zacks et al. 2001; Speer et al. 2003, 2007; DuBrow and Davachi 2013, 2016; Ezzyat and Davachi 2014; Baldassano et al. 2017; Ben-Yakov and Henson 2018).

Despite this degree of consistency across people in how they segment events, meaningful individual differences exist atop these shared tendencies. Individual differences in *behavioral* segmentation are stable over time (Speer et al. 2009), correlate with age and other cognitive abilities such as working memory capacity, long-term memory retrieval, and performance on other tasks (Zacks et al. 2006; Sargent et al. 2013; Bailey et al. 2017; Jafarpour et al. 2022), and are disrupted in certain clinical conditions such as schizophrenia (Zalla et al. 2004). Yet despite this ample behavioral evidence, investigations into individual differences in *neural* event segmentation have been very limited. This is due, in part, to the challenge of capturing individual boundaries (as opposed to relying on normative boundaries or using stimuli with pre-defined boundaries) while also preserving natural viewing (i.e., avoiding having subjects segment during encoding, which could alter the viewing experience and introduce unwanted effects on brain activity, or segment upon a second, biased viewing). However, recently developed algorithms for identifying latent brain state changes (Baldassano et al. 2017; Geerligs et al. 2021) allow us to infer neural event boundaries from fMRI data acquired during passively viewed continuous “naturalistic” stimuli (ex: movies) without prior knowledge of boundary locations.

Characterizing individual differences in neural event segmentation within a region during natural, passive viewing can extend our understanding of both cortical variability in event segmentation and the functional role of segmentation in cataloging experiences. Individual neural activity at normative event boundaries during a narrative reading task has been shown to predict the organization of information into long-term memory (Ezzyat and Davachi 2011). Here we sought to extend this work: above and beyond differences in objective recall accuracy, complex narrative stimuli generate personalized, idiosyncratic memories and appraisals (Black and Bower 1980), which could arise in part from individual differences in event segmentation during encoding—in other words, how the same (presented) events are (uniquely) segmented. We hypothesized that an individual’s pattern of neural event boundaries during movie-watching would predict how they ultimately remembered and appraised the stimulus. Further, these effects might be somewhat stimulus-dependent: different stimuli (in our case, movies) might evoke more or less individual variability overall, more or less individual variability in specific cortical regions, and/or differences in the specific regions where neural event boundaries relate to ultimate appraisal. We investigated these questions in a dataset of healthy adults who freely viewed and appraised four short movies during fMRI scanning.

To preview our findings, we first demonstrated that our chosen algorithm for automatically inferring neural event boundaries (Baldassano et al. 2017) can operate reliably at the individual level. Using individually defined neural boundaries, we found that across-subject variability followed a posterior-to-anterior cortical gradient: sensory processing regions were most consistent across individuals, while higher-order association regions were most variable. We found evidence for a relationship between individual event boundaries and the ultimate recall and appraisal of narratives in a subset of regions that also showed normative alignment with behavioral boundaries in our movies. The specific regions showing these relationships were different across movies, suggesting that depending on stimulus, distinct regions support aspects of online “chunking” that impact how the stimulus is later remembered and appraised.

## Materials and Methods

### Experiment 1 (main fMRI experiment)

We recruited and scanned a total of 48 subjects (all native English speakers; 27F, median age-24.5, range= (19,64)) at the National Institutes of Health (NIH). All subjects provided informed written consent prior to the start of the study in accordance with experimental procedures approved by the Institutional Review Board of the NIH. We discarded incomplete datasets without all four movies (described below), leaving 43 subjects whose data we analyzed here. Subjects were compensated $50 per hour for their time.

Subjects watched four movies (ranging from 7:27-12:27 min each) while we collected fMRI data using a 3T Siemens Prisma scanner with a 64-channel head coil. Movie order was pseudorandomized for each subject such that order was counterbalanced at the group level. Functional images were acquired using a T2*-weighted multiband, multi-echo echo-planar imaging (EPI) pulse sequence with the following parameters: TR = 1000 ms, echo times (TE) = [13.6, 31.86, 50.12 ms], flip angle = 60 deg, field of view = 216 x 216 mm, in-plane resolution = 3.0mm^2^, slice thickness 3.0mm, number of slices = 52 (whole-brain coverage), multiband acceleration factor = 4). Anatomical images were acquired using a T1-weighted MPRAGE pulse sequence with the following parameters: TR = 2530 ms, TE = 3.30 ms, flip angle = 7 deg, field of view = 256 x 256 mm, in-plane resolution = 1.0 mm^2^, slice thickness = 1.0 mm).

The movies were projected onto a rear-projection screen located in the magnet bore and viewed with an angled mirror. The experiment was presented using PsychoPy (Peirce et al., 2019). Following each movie (i.e., while still in the scanner) subjects completed a task battery designed to probe their interpretations and reactions to the movie, including the following: 1) a three-minute free recall/appraisal task in which subjects spoke freely about their memories and impressions of the movie, during which their speech was captured with a noise-canceling microphone; 2) multiple-choice comprehension questions designed to ensure they had been paying attention (ex: “At what holiday dinner does the father yell at the older son?” for *Growth*); 3) multiple-choice and Likert-style items assessing reactions to various characters and to the movie overall. See section “Measuring similarity in the recall” for the experimental instructions for the free recall/appraisal task and see study GitHub repository for all of the memory comprehension questions.

During a separate behavioral visit, subjects completed a battery of tasks and questionnaires including selected instruments from the NIH Cognition and Emotion toolboxes as well as other psychological and psychiatric self-report scales. These data are not analyzed or reported here.

### MRI data preprocessing

Following the conversion of the original DICOM images to NIFTI format, we used AFNI (Cox, 1996) to preprocess MRI data. Preprocessing included the following steps: despiking, head motion correction, affine alignment with subject-specific anatomical (T1-weighted) image, nonlinear alignment to a group MNI template (MNI152_2009), combination of data across the three echoes using AFNI’s “optimally combine” method, and smoothing with an isotropic full-width half-maximum of 4 mm. Each subject’s six motion time series, their derivatives, and linear polynomial baselines for each of the functional runs were included as regressors. All analyses were conducted in volume space and projected to the surface for visualization purposes.

We used mean framewise displacement (MFD), a per-subject summary metric, to assess the amount of head motion in the sample. MFD was overall relatively low (*Iteration* - mean=0.08, s.d. = 0.03, range = (0.03, 0.18), *Defeat*- mean=0.08, s.d. = 0.03, range = (0.04, 0.19), *Growth* - mean=0.07, s.d. = 0.03, range = (0.04, 0.17), *Lemonade* - mean=0.08, s.d. = 0.03, range = (0.04, 0.17) and did not differ across movies (repeated-measures ANOVA, F(3,126) = 1.44, *p*=.23).

We included an additional preprocessing step using a shared response model (Chen et al. 2015) in *BrainIAK* (Kumar et al. 2020) to account for different functional topographies across individuals. This step was performed for each of the four movies separately. First, we fit a model to capture the reliable whole-brain responses to the movie across subjects in a lower dimensional feature space (features = 50). We then applied this model to reconstruct the individual voxelwise time courses for each participant. This procedure serves as an additional denoising step and makes spatiotemporal patterns more consistent across subjects.

### Experiment 2 (auxiliary behavioral experiment)

We collected an auxiliary behavioral dataset (i.e., outside the MRI scanner) at Dartmouth College to further refine and support results from *Experiment 1*. 44 subjects (21F, median age 20, range=18-32) were presented with the same paradigm as in *Experiment 1,* except that while watching each film, individuals were instructed to press a button each time they perceived that a new scene is starting (i.e., points in the movie when there is a “major change in topic, location, time, etc.”) and to expect scenes to be between 10 seconds and 3 minutes long. We used the same instructions used in Baldassano *et al.,* (2017). We discarded subjects who did not complete all four movies, leaving n=40 for our analysis. The paradigm was hosted and presented using JsPsych (de Leeuw 2015). Subjects were compensated $15 an hour for their time or given participation credit, and the study was approved by the Institutional Review Board of Dartmouth College.

### Movie overview

Our stimuli were four short films made by independent filmmakers that were chosen because they were rich and engaging, yet they depicted ambiguous scenarios that provoked divergent reactions and interpretations in different individuals. Three of the movies were “social” in nature and followed different narratives of humans taking actions and interacting, while the fourth depicted purely mechanical information (a long, complex Rube Goldberg machine that traversed a house). We chose independent films so that subjects would be less likely to have experienced the material before. In a debriefing questionnaire, three subjects reported having seen one of the movies (*Growth*) prior to the experiment.

Here we provide brief descriptions of each movie along with YouTube links. (Note that the versions presented to subjects were edited to remove credits and title pages; these edited versions are available upon request.) *Iteration* (https://youtu.be/c53fGdK84rc; 12:27 min:sec) is a sci-fi movie that follows a female character as she goes through multiple iterations of waking up and trying to escape a facility. A male character appears towards the end to help her. *Defeat* (https://youtu.be/6yN9VH_4GSQ; 7:57 min:sec) follows a family of three (mother, two children) as the brother bullies his sister and she builds a time machine to go back and get revenge. *Growth* (https://youtu.be/JyvFXBA3O8o; 8:27 min:sec) follows a family of four (mother, father, two brothers) as the children grow up and eventually move out amid some family conflict. *Lemonade* (https://youtu.be/Av07QiqmsoA; 7:27 min:sec) is a Rube-Goldberg machine consisting of a series of objects that move throughout a house and ends in the pouring of a cup of lemonade. This movie was lightly edited to remove fleeting shots of human characters. *Iteration* and *Defeat* both contained screen cuts (continuity editing), while *Growth* and *Lemonade* were shot in a continuous fashion with the camera panning smoothly from one scene to the next.

### Automatic event boundary detection

To automatically identify neural event boundaries from fMRI data, we fit a series of Hidden Markov Models (HMMs) (split-merge option; Baldassano et al., 2017) as implemented in the *BrainIAK* toolbox (Kumar et al. 2020), adapting code made available on the *Naturalistic Data* tutorial website (Chang et al. 2020). The HMM approach does not rely on annotations or hand-demarcated events, but rather infers event boundaries from shifts in stable patterns of brain activity. It relies on voxelwise patterns *within* regions. We restricted our analyses to the neocortex and used the 100-parcel, 7-network Schaefer parcellation (Schaefer et al. 2018), in keeping with past work using HMMs to identify event boundaries (Cohen and Baldassano 2021).

The HMM approach assumes that each event has a distinct signature of activity that shifts at event boundaries within region. Specifically, the model assumes that (1) each subject starts in an event and then each forthcoming timepoint is either in the same event (state) or in the next state, and (2) that the voxelwise pattern of activity in a region is correlated across timepoints within the same event. The model identifies both the optimal number of events and then the transitions between these events. We were ultimately interested in individual variability in the location of event boundaries. While we acknowledge that there is likely interesting individual variability in the number of events (i.e., segmentation rate) in each region, this is a somewhat different question than variability in the location of neural event boundaries: consider that even if two individuals have the same number of events in a region, the specific locations of their event boundaries could still be completely different. The location of boundaries is more directly related to the movie content onscreen at any given moment than the overall number of events, and therefore of greater scientific interest to us here, given that our goals were to investigate 1) how moment-to-moment segmentation patterns relate to ultimate appraisals, and 2) how movie content affects the locations and degree of this variability. In addition to this theoretical justification, there are two related methodological justifications to fixing the number of events within a region while allowing locations to vary across subjects. First, we are using an inherently noisy method to infer neural event boundaries which was created to work well at the group level. Therefore, by fixing the number of events using group-average data, we constrain the individual HMM solutions to a reasonable number of events. Second, it is not clear how to reliably estimate the optimal number of events at the individual level, since training and test data need to be independent.

Therefore, to focus on individual variability in boundary locations, we first determined the optimal number of events (*k*) for each region at the group level using a train-test split procedure (with different subjects in the train and test groups). We used a fairly liberal range of possible values for *k*: the minimum allowed *k* was 6, and the maximum (90-150) differed slightly across movies owing to their different lengths, but always reflected a minimum average event length of five seconds. For each movie and each region, we split the subjects into train/test groups (each with 21/22 subjects) and averaged voxelwise timecourses in each group, resulting in one average voxel-by-time array each for the training and test data. We then fit a series of models to the average training data using each possible value of *k* and tested each model on the average test data, assessing model fit using the log-likelihood. Having repeated this procedure for each *k*, we took the *k* value with the maximum log-likelihood as the optimal number of events within this region for this train/test split. We repeated this procedure 100 times with different train/test splits within each region. For each region, we then calculated the median optimal *k* value across these 100 iterations and used this as the *k* value for our subsequent analyses.

Using these fixed values of *k,* for each region, subject, and movie, we then fit an HMM to that subject’s individual timeseries to identify the location of implicit event boundaries. Therefore, we had one set of boundary locations for each region for each subject for each movie.

Notably, the fit of the HMM (as measured by the log-likelihood) is tightly coupled with the number of events and our movies do differ in length (such that the model fit is best with the shortest movie and worst with the longest movie). When accounting for the difference in movie length, the model fit does not differ across movies. Model fit was also highest in sensory regions (ex: V1) and decreased on a cortical gradient in multimodal regions; this illustrates that regions where there is more consistency across people (c.f. Figure 1), are also regions where the model is performing better, implying that more shared boundaries are not due to poor model performance.

**Figure 1.**
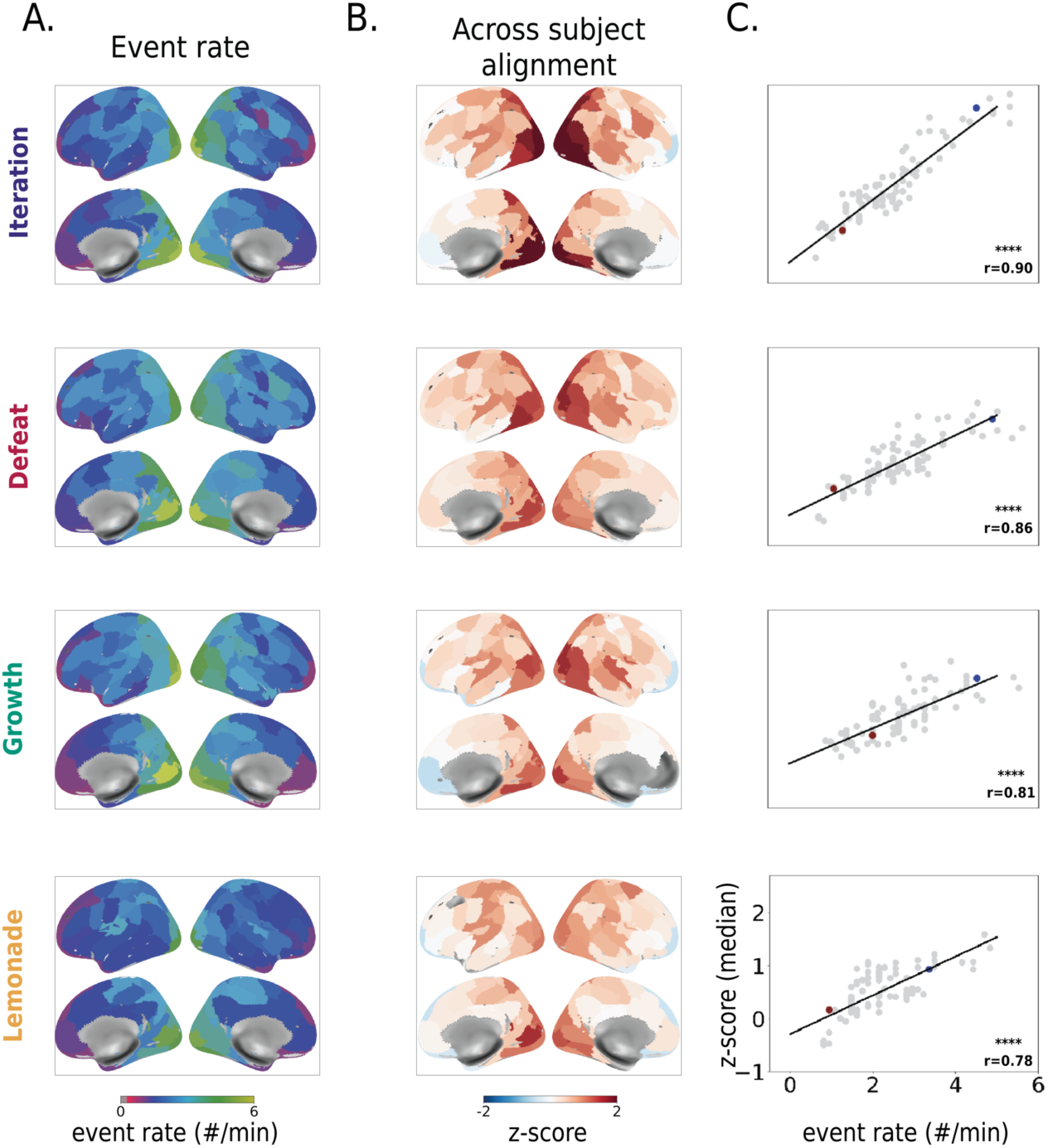
Segmentation rate and degree of idiosyncrasy in boundary locations covary across the cortex. **A. Event rate.** The optimal number of events for each region (determined at the group level) follows the expected cortical gradient: faster segmentation (more events) in posterior, sensory regions and slower segmentation (fewer events) in anterior, higher-order regions. **B. Across-subject alignment within movie.** There is generally above-chance alignment in the location of event boundaries among individuals across the cortex (*n*=10,000 bootstraps, *q*<.05, FDR corrected across 100 regions). Alignment is highest in posterior sensory regions and lowest in anterior association regions. The degree of alignment in each region varies slightly across movies. While greater-than-chance alignment was found in the majority of regions (>89/100) in each movie, regions that did not show significant alignment tended to be in higher-order regions such as the prefrontal cortex. **C. Correlation between event rate and within-movie alignment.** In all four movies, event rate and degree of alignment were strongly correlated across regions (Spearman *r* = .78-.90, *p*<.001) such that fast transitioning regions were more closely aligned across subjects, while slower regions were less aligned (i.e, showed more individual variability). Each gray dot indicates a region; we highlight a visual region in blue and a prefrontal region in red. Legend: **** indicates *p*<.0001.

### Computing alignment across subjects

We used permutation testing to quantitatively assess the consistency of boundary locations across individuals. For each movie, region, and pair of subjects, we permuted boundary locations n = 1000 times to derive a z-score for the match between true boundaries relative to the null distribution. In both the true and permuted data, boundaries that were within 3 TRs of one another were counted as a match, consistent with past work (Baldassano et al. 2017; Williams et al. 2022). We then created subject-by-subject matrices of these z-scores in every region. Importantly, depending on which member of a pair of subjects was permuted (versus treated as the “ground truth”), there were slight differences in the resulting z-score; we took the mean of the upper and lower triangle of these matrices for relevant subsequent analyses.

### Controlling for possible confounds

Several factors outside the ongoing cognitive processes of interest could contribute to higher or lower alignment in detected boundary locations between a given pair of subjects. The factors detailed below were used as regressors of no interest to control for these unwanted influences in the ensuing analyses, as described in subsequent sections.

### Inter-subject correlation (ISC) of head motion

It is possible that shared head motion at similar moments in the movie could lead the HMM to (perhaps falsely) detect similar neural event boundaries in a given pair of subjects. To control for this possibility, we computed the inter-subject correlation (ISC) of the framewise displacement across time for each subject pair. We then used this subject-by-subject motion-ISC matrix as a nuisance regressor in subsequent analyses.

### Overall head motion

Similarly, subjects with high overall levels of head motion likely have lower-quality fMRI data, which could bias the detection of neural event boundaries altogether (though importantly, as detailed in “MRI data preprocessing”, absolute levels of head motion were relatively low in our sample). To control for this possible confound, we used each subject’s median framewise displacement (FD) across all timepoints and generated a subject-by-subject similarity matrix using the Anna Karenina principle (“all low (or high) motion scorers are alike; each high (or low) motion scorers is different in their own way”) by taking the mean score between each subject pair (Finn et al. 2020).

### Memory performance

Some subjects may simply have been paying better attention throughout the scan session and/or during certain movies, which could generate stronger stimulus-driven neural responses and in turn drive up similarity in event boundaries among these subjects. We controlled for this possibility using subjects’ performance on the four multiple-choice memory recall questions presented at the end of each movie, which were designed to be quite difficult. We computed each subject’s memory performance score as the fraction of correct responses on these questions, and generated a subject-by-subject similarity matrix of these scores also according to the aforementioned Anna Karenina principle (“all low (or high) memory scorers are alike; each high (or low) scorer is different in their own way”) by taking the mean score between each subject pair (Finn et al. 2020). The same approach was applied to the behavioral boundaries collected in Experiment 2.

### Across-subject alignment within movies

The goal of this analysis was to assess the degree to which event boundary locations were aligned across subjects within each movie. Subject-by-subject matrices of alignment values (z-scores) in every region, generated using the permutation method described above (see section “Computing alignment across subjects”), were used.

### Experiment 1

We fit a linear regression to regress out (1) head-motion ISC, (2) overall head motion, and (3) memory performance (See “Controlling for possible confounds” for more detail) from the subject-by-subject boundary alignment matrix in each region. Using the residuals of this regression, we took the median alignment value (z-score) across subject pairs as the summary statistic for each region. To determine whether alignment was significantly above chance (i.e., greater than 0), we performed this same calculation in a subject-wise bootstrapping framework (*n*=10,000 bootstraps) to create a non-parametric null distribution and compared our observed median residual z-score to this distribution to calculate a p-value. P-values were corrected for multiple comparisons using the false discovery rate (FDR) based on the number of regions in our parcellation (100) using an alpha of .05.

### Experiment 2

We fit a linear regression to regress out memory performance (See “Controlling for possible confounds” for more detail) from the subject-by-subject matrix of alignment in behaviorally reported event boundaries. Using the residuals of this regression, we took the median alignment value (z-score) across subject pairs as the summary statistic. To determine whether alignment was significantly above chance (i.e., greater than 0), we performed this same calculation in a subject-wise bootstrapping framework (*n*=10,000 bootstraps) to create a non-parametric null distribution and compared our observed median residual z-score to this distribution to calculate a p-value.

### Measuring cross-subject similarity in movie recall/appraisal

We used an inter-subject representational similarity analysis (IS-RSA) approach (Glerean et al. 2016; Chen et al. 2020; Finn et al. 2020) to investigate whether pairs of subjects that were similar in their neural event boundaries in a region also had similar interpretations of the movie.

To elicit appraisals, we presented the following prompt to subjects immediately following each movie: “During this section, you will have three minutes to say what you remember about the video. You can talk about characters, events, your opinions, or anything else that comes to mind. Try to fill the whole three minutes once the timer appears and remember-there are no wrong answers!” Subjects spoke freely while their speech was recorded using a noise-canceling microphone. These recordings were professionally transcribed and minimally cleaned to remove interjections, such as “um” or “uh”, and repeated words. One subject’s recall data was not able to be transcribed due to being corrupted by scanner noise and was discarded from this analysis, leaving n=42.

The total number of words spoken varied across subjects and movies: *Iteration*: - mean=386, s.d. = 80, range = 213-547, *Defeat* - mean=362, s.d. = 78, range = 153-533, *Growth* - mean=368, s.d. = 72, range = 243-520, *Lemonade* - mean=359, s.d. = 84, range = 180-576. The number of words differed significantly between movies (repeated-measures ANOVA, F(3,123) = 2.87, *p*=.04). Post hoc tests showed that the number of words used in *Iteration* was significantly greater than the number of words used in *Defeat* (paired *t*-test, *t*=2.81, *p*=.01) and *Lemonade* (paired *t*-test, *t*=2.12, *p*=.04), which is not surprising given that *Iteration* was the longest movie (∼12 minutes as compared to ∼8 minutes for the other 3 movies). Since our goal in this analysis was to compare subject-to-subject similarity in boundary location to similarity in appraisal *within* a movie as opposed to *across* movies, differences in the number of words used across movies should not influence our results in a meaningful way. Importantly, the degree of similarity in the appraisal of *Iteration* is not significantly higher compared to the other movies. In fact, the highest similarity was in *Growth*, which was greater than *Iteration*, *Defeat*, and *Lemonade* (paired *t*- tests, *t*≥11.84, *p*<.001) (median similarity - *Iteration* - 0.58, *Defeat*- 0.59, *Growth*-0.63, and *Lemonade* - 0.58; repeated-measures ANOVA, F(3,2580) = 86.06, *p*<.001). Therefore, there does not seem to be a direct relationship between the number of words used to appraise a movie and the degree of similarity in appraisals across subjects.

The text from each individual’s speech data was then embedded into a 512-dimensional space via Google’s Universal Sentence Encoder (USE; (Cer et al. 2018); implemented via https://www.tensorflow.org/hub/tutorials/semantic_similarity_with_tf_hub_universal_encoder). We chose the USE model in part because it was trained to identify similarities between pairs of sentences, and, as a sanity check, because it was best able to differentiate appraisals from different movies (i.e., recall between *Growth*-*Growth* was more similar than recall between *Growth*-*Defeat*, etc.) whereas this sensitivity was absent or weaker with other pre-trained context-sensitive models that we tested (including BERT_BASE_, MiniLM and MPNet (Devlin et al. 2019; Song et al. 2020; Wang et al. 2020)). USE was also used on event descriptions in a recent publication (Lee and Chen 2022). We computed the cosine similarity between the 512-dimensional vectors generated by USE to measure the semantic similarity between pairs of subjects in their interpretations, resulting in one subject-by-subject appraisal similarity matrix per movie. In an IS-RSA, we then used a Spearman correlation to compare the lower triangle of these appraisal similarity matrices to the lower triangle of the boundary alignment matrices in each region. A partial Mantel test (nPerms = 10,000; *q*<.05, FDR corrected across regions) was used to assess the relationship between event boundary alignment and appraisal similarity while controlling for memory performance (See “Controlling for possible confounds” for more detail). By controlling for memory performance, we were able to bias our findings toward relationships driven more by subjective impressions and interpretations than objective recall *per se*.

### Identifying stimulus-dependent properties by taking the difference in alignment across movies

The goal of this analysis was to assess the degree to which stimulus content influences across-subject alignment in event boundary locations by comparing subject-by-subject alignment matrices between pairs of movies.

### Experiment 1

For a given region and pair of movies, we fit a linear regression to model the difference between the two subject-by-subject alignment matrices as a function of the following regressors of no interest: (1) difference in head-motion ISC, (2) the difference in overall head motion, and (3) the difference in memory performance between the movies (See “Controlling for possible confounds’’ for more detail). We then took the median value from this residual difference matrix and compared it to a non-parametric null distribution (n = 10,000 bootstraps) to calculate a p-value. P-values were corrected for multiple comparisons using the false discovery rate (FDR) based on the number of regions in our parcellation (100) using an alpha of .05. To investigate the overall (i.e., whole-brain) variation in alignment across movies, we employed a linear mixed-effects model, implemented using lme4 and lmerTest (Bates et al. 2015) in R. Estimated marginal means were calculated using the emmeans package (Lenth 2023). We used the within-movie alignment regression’s intercept (accounting for the regressors of no interest, see *Methods* section - “Across-subject alignment within movies”) by movie, treating region as a random effect.

### Experiment 2

For a given pair of movies, we fit a linear regression to model the difference between the two subject-by-subject alignment matrices as a function of the following regressor of no interest: the difference in memory performance between the movies (See “Controlling for possible confounds” for more detail). We then took the median value from this residual difference matrix and compared it to a non-parametric null distribution (*n* = 10,000 bootstraps) to extract a p-value. To investigate the variation in behavioral alignment across movies, we adopted the method used by Chen et al. (2017) and fit a linear mixed-effects model predicting alignment by movie and memory (our regressor of no interest) with crossed random effects. This approach allowed us to account for non-independence of the pairwise alignment in the data from repeated observations for each participant. To account for redundancy, we manually adjusted the degrees of freedom and standard error, as suggested by Chen et al. (2017).

### Comparison of normative neural event boundaries and normative behavioral event boundaries

We used data from *Experiment 2* to identify a set of group-level or “normative” behavioral boundaries for each movie. Individuals’ button presses were first rounded to the nearest second. To make the behavioral boundaries commensurate with the HMM-derived neural boundaries, we needed to account for the delay in the hemodynamic response, which may be partially offset by the delay in behavioral motor responses (i.e., button presses) after a boundary is detected. Therefore, we added 3 TRs/seconds to each button press (without allowing for events past the end of the movie).

We took two approaches, a density-based and a peak-based method (described in turn below), to compute the similarity between normative HMM-derived boundaries and behavioral-study normative boundaries.

### Density Alignment Method

We generated a density distribution of “button presses” to indicate, at each timepoint (second), what percentage of subjects in the behavioral study marked a boundary location at that timepoint (or within +/- 3 s of that timepoint). Specifically, a binary distribution (event or no event) over time was generated for each subject. These subject-level distributions were combined to get a density distribution reflecting the proportion of subjects indicating an event per timepoint. We then computed a Pearson correlation between this behavioral density distribution and the density distribution of individual-subject HMM-derived boundaries for each region and compared this “true” correlation to a null distribution of correlations that were generated by “rolling” (or circle-shifting) the HMM-derived boundaries at each TR (thus the number of permutations was limited to the number of TRs for each movie). This generation of a proportion-based density distribution is similar to the “segmentation agreement” previously used to compare individuals to the group average (Zacks et al. 2006; Bailey, Kurby, et al. 2013; Bailey, Zacks, et al. 2013).

### Peak Alignment Method

We used our permutation-based alignment metric (See “Computing alignment across subjects”), treating the brain as the “ground-truth” (i.e. permuting behavioral boundaries) to compare the behaviorally-derived boundaries to the group-average HMM-derived boundaries for each region. Normative (group-average) HMM-derived boundaries were computed by averaging voxelwise activity across subjects and then fitting an HMM to data within each region using the pre-determined number of events for that region. To determine the normative boundaries from the behavioral study for this approach, we needed “peak” shared behavioral boundaries. For this, we defined the peaks as timepoints when >50%, or at least 21/40, subjects marked a boundary and enforced local sparsity by limiting peaks to timepoints that were more than 3 seconds apart. (If two peaks were within 3 seconds of one another, we took the higher “peak” [more agreement]; if they were of equal height, we took the median timepoint.) Importantly, we used 3 seconds as our tolerance for small deviations in button-press times across subjects to be in line with the fMRI data, where we had used 3 TRs (= 3 s).

### Combining Methods

To declare significant alignment between normative behavioral and normative neural boundaries in a region, that region had to show above-chance alignment with behavioral boundaries (*p*<.05) in both the density and peak methods.

### Code Accessibility

Data analysis, including links to code and other supporting material, can be found at: https://github.com/thefinnlab/individual_event_seg/.

### Data Accessibility

Data from this study, including raw MRI data, will be made available on OpenNeuro upon publication. Other data including the behaviorally reported boundaries and the full transcribed recall and appraisal (text) can be found at: https://github.com/thefinnlab/individual_event_seg/tree/main/data.

## Results

In this work, our main goal was to investigate variability in neural event segmentation at the individual level and its behavioral consequences. We automatically detected region-wise event boundaries in individual subjects and used these to quantify the degree of variability across the cortex and how this variability changes with stimulus content. We also show how and where similarity between individuals’ neural event boundaries during encoding can predict similarity of recall and interpretation, thereby highlighting a functional role for event segmentation in an individual’s ultimate appraisal of an experience.

### Slower-segmenting regions show more individual variability

Forty-three subjects watched four movies during fMRI scanning. These differed in meaningful and important ways (see *Methods* - “Movie Overview” and *Results* - “Degree of across-subject alignment varies with movie content.”). While we hypothesized that these differences would impact individual variability (an idea we return to in the following section), our goal with this first analysis was to demonstrate that cortical patterns of event segmentation, and individual variability therein, are generally consistent irrespective of the stimulus used.

To define neural event boundaries, we used a recently developed algorithm for detecting event boundaries in fMRI data: the Hidden Markov Model (HMM) first proposed by Baldassano et al. (2017), which does not rely on annotations or hand-demarcated events but rather detects event boundaries as shifts in stable patterns of brain activity within a region. While this algorithm yields valid neural event boundaries at the group level, to our knowledge, it had not been previously applied to individual-subject data. Therefore, we first undertook an analysis to assess whether this model can stably and reliably detect event boundaries at the individual level. All fMRI data were parcellated into 100 cortical regions using the Schaefer atlas (Schaefer et al. 2018).

We first sought to replicate past work (Baldassano et al. 2017; Geerligs et al. 2022) showing that at the group level, the number of events (i.e., the granularity of segmentation) is higher in sensory regions and lower in higher-order association areas that are sensitive to narrative information at longer timescales (Lerner et al. 2011; Honey et al. 2012). Using a train-test procedure to determine the optimal number of events (*k*) for each region from group-average data, we demonstrate that this relationship is maintained irrespective of the stimulus: event rate (number of events per minute) follows a posterior-to-anterior gradient (**Fig. 1A**).

We then investigated to what degree segmentation varied across individuals. Event segmentation can vary both at the level of the number of events (*k*) and the location of boundaries; we focus here on the latter, both for methodological reasons and because it is more directly related to movie content (see *Methods* section “Automatic event boundary detection” for more details). We used the fixed value of *k* for each region determined at the group level (cf. **Fig. 1A**) to fit an HMM to each individual subjects’ data from that region (see *Methods* section “Automatic event boundary detection”). We then assessed to what degree event boundary locations were aligned across subjects using a permutation-based method that generates a z-score for observed alignment relative to an appropriate null distribution for the total number of boundaries in each region (see *Methods* sections “Computing alignment across subjects” and “Across-subject alignment within movies-Experiment 1”).

Across all four movies, event boundaries were significantly aligned across subjects in most regions of the brain (>89/100), but this degree of alignment varied on another posterior to anterior gradient such that event boundary locations were more shared in sensory regions and more idiosyncratic in higher-order regions (**Fig. 1B**). We established strong correlations (*r*= .78-.90) between event rate and the degree of alignment across all four movies: faster-segmenting regions showed higher alignment across subjects (less individual variability), while slower-segmenting regions showed less alignment across subjects (more individual variability; **Fig. 1C**).

Notably, insignificant or negative across-subject alignment was seen in 20 unique regions across the four movies (1 for *Defeat,* 9 for *Iteration,* 11 for *Growth,* and 10 for *Lemonade*; negative across-subject alignment would indicate that permuted boundaries have a higher chance of showing alignment than the true boundaries). These regions were located in the prefrontal cortex (14/20), orbitofrontal cortex (2/20), temporal pole (3/20), and the cingulate (1/20). Of these, the majority were higher order “default mode” (40%) or limbic regions (25%) that have the slowest timescales of information processing (Geerligs et al., 2022; Hasson et al., 2008) and show more idiosyncratic anatomy and function (Hill et al. 2010; Mueller et al. 2013), including during naturalistic stimulation (Vanderwal et al. 2017; Finn et al. 2020; Gao et al. 2020; Chang et al. 2021). (It should also be noted, however, that these regions are also most commonly affected by signal drop-out.) No parcel showed negative or insignificant alignment across all four movies. (All analyses in this section were controlled for the effects of head motion and overall memory performance; see *Methods* section “Controlling for possible confounds”).

### Degree of across-subject alignment varies with movie content

Having established that there is significant across-subject alignment that follows a cortical gradient in all movies, we next aimed to quantify how the degree of individual variability in event segmentation within region differed across the four movies. To help validate any differences across movies in neural event segmentation, we conducted an auxiliary behavioral experiment (Experiment 2 - see *Methods*) in which a separate set of 40 subjects at Dartmouth College performed the same task as Experiment 1 outside the scanner, except that while watching each movie, individuals were instructed to press a button each time they thought there was an event boundary; i.e., points in the movie when there is a major change in topic, location, time, etc. This allowed us to investigate whether similar patterns of movie-dependent variability in neural event segmentation were also present with behavioral event segmentation.

The movies differed in numerous and important ways. For the purposes of this project, we focus on two hypothesized dimensions: (1) the presence or absence of screen cuts, and (2) the presence or absence of social and affective information. Three of our movies followed a character-driven narrative trajectory complete with social and affective information (*Iteration, Defeat,* and *Growth*), while the fourth (*Lemonade*) followed a series of mechanical events (Rube-Goldberg machine) as opposed to a human-driven plot line. Two—*Iteration* and *Defeat*—used continuity editing (i.e., screen cuts), which serve as a strong cue for the start of a new event (Schwan et al. 2000; Zacks et al. 2010; Magliano and Zacks 2011; Smith et al. 2012; Loschky et al. 2020). We hypothesized that both dimensions would facilitate segmentation and increase the degree of across-subject alignment. First, continuity editing (such as screen cuts) is a well-documented driver of event boundaries; it often indicates a scene change and is highly correlated with situational changes that indicate the start of a new event (Schwan et al. 2000; Zacks et al. 2010; Magliano and Zacks 2011; Smith et al. 2012; Loschky et al. 2020). Second, socio-affective content often engages shared schemas of social interactions (ex: family dinners in *Growth*; (Dunbar 1998; Wood et al. 2003; Adolphs et al. 2016) and is often more engaging than its non-social counterparts. Both social and affective information induce shared neural responses during movie-watching (Finn et al. 2018; Chen et al. 2020; Song et al. 2021) and recall (Chen, Leong, et al. 2017; Tomita et al. 2021) and may aid in segmentation (Boggia and Ristic 2015; Ristic and Capozzi 2022). It is notable, however, that, paradoxically, along with these shared responses, such dynamic, emotional content is also better suited at pulling out nuanced individual differences (Finn and Bandettini 2021), which we explore in the next section.

In our dataset, *Iteration,* a social movie with screen cuts, had significantly higher self-reported arousal (which we take as a proxy for affect/emotion) than the other three movies (repeated-measures ANOVA, F(3,126)=7.49, *p*=.0001; *Iteration > Defeat, Growth, Lemonade; t* ≥ 3.68, *p*<.001). Overall, self-reported engagement levels did not differ between movies (repeated-measures ANOVA, F (3,126) =1.67, *p*=.18), although in pairwise comparisons, *Iteration* had mildly higher self-reported engagement levels than *Defeat* (pairwise comparisons, *t*=2.2, *p*=.03). Thus, we expected that we would see the following rank-order from highest to lowest in across-subject consistency: (1) *Iteration* (social information, screen cuts and most emotionally arousing), (2) *Defeat* (social information and screen cuts), (3) *Growth* (social information, but no screen-cuts), and (4) *Lemonade* (no social-content and no screen cuts).

We tested these hypotheses by comparing alignment in each cortical region between each pair of movies (see *Methods section –* “Identifying stimulus-dependent properties by taking the difference in alignment across movies.”) (All analyses in this section were controlled for effects of head motion [in neural data] and overall memory performance [in both behavioral and neural data]; see *Methods* section “Controlling for possible confounds”.) We used a linear mixed-effects model to assess the effect of movie on across-subject alignment, with region treated as a random effect. The analysis revealed a significant effect of movie (F(3, 300) = 39.47, *p*<.001). Post-hoc tests were conducted to compare the estimated marginal means (EMMs) of the four movies. The results showed that alignment in *Iteration* was significantly higher than *Defeat* (estimate = 0.09, *p*=.029), *Growth* (estimate = 0.27, *p*<.001), and *Lemonade* (estimate = 0.29, *p*<.001). Additionally, alignment in *Defeat* was significantly higher than Growth (estimate = 0.18, *p*<.001) and *Lemonade* (estimate = 0.20, *p*<.001), while *Growth* and *Lemonade* did not differ significantly (estimate = 0.02, *p*=.887). Thus, we observed the following rank order of movies, from highest to lowest neural alignment: [*Iteration*] > [*Defeat*] > [*Growth* and *Lemonade*] (**Fig. 2A**). Critically, variability in behavioral alignment largely mirrored this rank order of movies: while we found significant overall alignment in behavioral event boundaries within all movies (*p*=.0001; nbootstraps=10,000-see *Methods* section “Across-subject alignment within movies.- Experiment 2”), we further observed that movies that evoked more variability in neural event segmentation (across all brain regions) also tended to evoke more variability in behavioral event segmentation ([*Iteration*] > [*Defeat*]> [Growth] > [Lemonade]; linear mixed effects modeling with movie as fixed effect; pairwise comparisons of the estimated marginal means are shown in **Fig. 2B** and bootstrapped pairwise comparisons are shown in **Fig. 2C**, Behavioral (right) columns).

**Figure 2.**
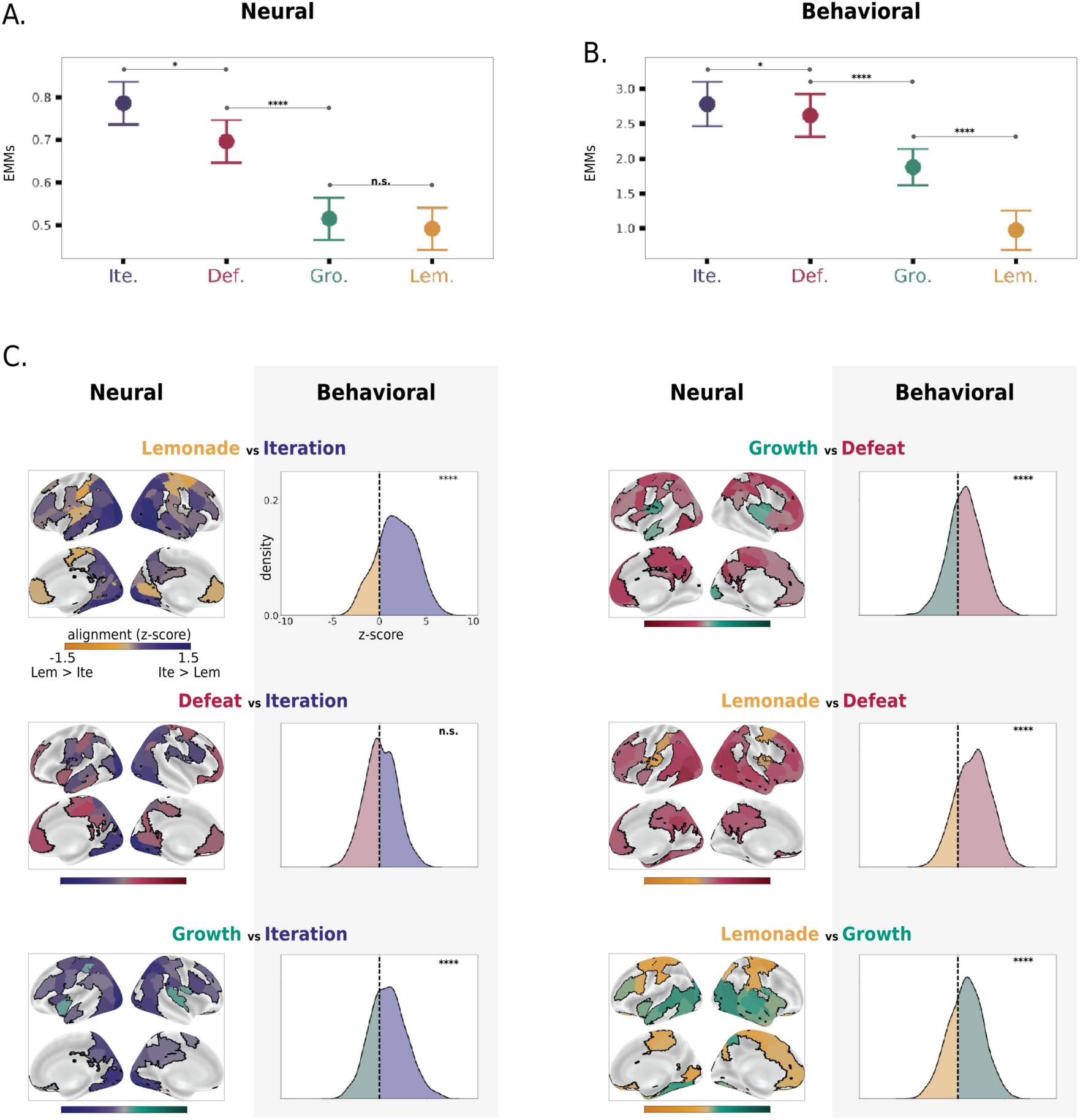
Degree of individual variability in neural and behavioral event segmentation depends on stimulus content. **A. Degree of overall alignment in neural event boundaries differs by movie.** Plots depict the distribution of the estimated marginal means (EMMs) for the alignment across all regions (*n* = 100) for each movie. Whole-brain alignment shows the following rank order across movies: [*Iteration*] > [*Defeat*] > [*Growth* and *Lemonade*]. **B. Degree of alignment in behavioral event boundaries differs by movie.** Plots depict the distribution of marginal means for the behavioral alignment across subjects. Across-subject behavioral alignment shows a rank order across movies that largely corresponds to the rank order of neural alignment: [*Iteration*] > [*Defeat*] > [*Growth*] >[*Lemonade*]. **C. Cortical variability in degree of alignment across movies and comparison with degree of behavioral alignment.** Left Columns-Pairwise comparisons between movies indicate regions where neural alignment is significantly higher in one movie versus another. Coloring indicates the movie where there is higher alignment and shade reflects the magnitude of the difference (maps are thresholded at *q*<.05; FDR corrected to 100 regions within movie pair). Significance was determined using the residuals after regressing the effects of head motion and memory performance. ***Right Columns-*** Pairwise comparisons between movies indicate cases where behavioral alignment is significantly higher in one movie versus another. For visualization purposes, we are showing the distribution of the difference in alignment values “z-scores” between pairs of subjects (the movie on the left was subtracted from the movie on the right). Significance was determined using the residuals after regressing the effects of memory performance (bootstraps=10,000). Note that the dominant directionality of each pairwise comparison (i.e, which movie shows higher alignment) is similar in the neural and behavioral data. Legend: * indicates *p*<.05; **** indicates *p*<.0001.

Again, these four movies differ along many dimensions that could influence the degree of individual variability in boundary locations. To formally assess whether our two *a priori* hypothesized features account for significant variance in across-subject alignment, we modeled these features (instead of movie identity itself) as fixed effects in a series of follow-up analyses. For the neural data, we ran two independent linear mixed effects models: one with social information (present/absent) and one with continuity editing (present/absent) as the fixed effect. Each model contained region as the random effect. Both social information and continuity editing were significant predictors of neural alignment in their respective models (social: beta - .17, p<.0001; continuity editing: beta - .24, p<.0001). However, running a single model that included both social information and continuity editing revealed that when considered jointly, only continuity editing was a significant predictor of alignment (beta-.23, p<.0001). For the behavioral data, both social information and continuity editing were significant predictors of behavioral alignment when treated as fixed effects in a single model (social: beta - .91, p<.0001; continuity editing: beta - .80, p<.0001) that also included subject-level crossed-random effects and memory scores as a fixed effect.

Thus, consistent with our hypotheses, we found that both behavioral and neural alignment were higher in movies in which there was a human-driven, social, and emotionally-engaging narrative trajectory, especially when there were screen cuts cueing new scenes (*Iteration*, *Defeat*). In movies where there were no screen cuts (*Growth* and *Lemonade*), across-subject alignment was lower. This supports the inference that segmentation is more straightforward for certain types of stimuli irrespective of the modality (neural/HMM-derived or behaviorally-derived).

Next, we took a region-specific approach to determine where in the cortex showed the biggest differences in alignment between movies (in other words, to identify brain regions where the degree of alignment is most content-dependent). Results are shown in **Fig. 2C** (Neural (Left) Columns; nbootstraps=10,000). We identified a trend such that certain movie content (social information) drives more alignment in neural event boundaries in mid-level regions of association-cortex. For instance, while regions on either end of the cortical hierarchy showed consistently low (PFC) or high (V1) alignment across movies (cf. Fig. 1B), canonical social-processing regions such as the superior temporal lobe (STL) showed higher alignment in movies with social interactions compared to non-social movies ([*Iteration*, *Defeat*, *Growth*] > [*Lemonade*]). Our non-social movie, *Lemonade,* showed primarily higher alignment in somatomotor processing regions, likely reflecting neural state changes in response to the continuous switches in the location and object-driven motion that are specific to this movie.

### Shared neural event boundaries lead to shared interpretations

We next tested the hypothesis that individual variability in neural event segmentation would have behavioral consequences, such that individuals who were more similar in their neural event boundaries during encoding would also be more similar in their interpretation of the movie. To do so, we used free-speech data acquired immediately following each of the four movies in which subjects were prompted to speak for three minutes about what they remembered and how they felt about the events and characters (see *Methods* section “Measuring cross-subject similarity in movie recall/appraisal” for exact instructions). Crucially, subjects did not simply recall the events objectively as they might do in a typical episodic memory task, but also shared subjective interpretations and overall reflections on the movies, often including links to their own personal lives. Thus, we henceforth refer to this task as the appraisal task.

We recorded and transcribed each subject’s speech and submitted the transcripts to Google’s Universal Sentence Encoder (USE), a tool from natural language processing (NLP) that encodes text into high-dimensional vectors that reflect semantic content (Cer et al. (2018); see *Methods* section “Measuring cross-subject similarity in movie recall/appraisal”). Language embeddings provide a relatively unbiased way to quantify similarity in appraisal content. For each movie, we then calculated a subject-by-subject similarity matrix from these vectors and compared it to the subject-by-subject similarity matrix of neural event boundaries in each region while controlling for objective memory performance (as measured by performance on memory questions) using an inter-subject representational similarity analysis (**Fig. 3A** - inset).

**Figure 3.**
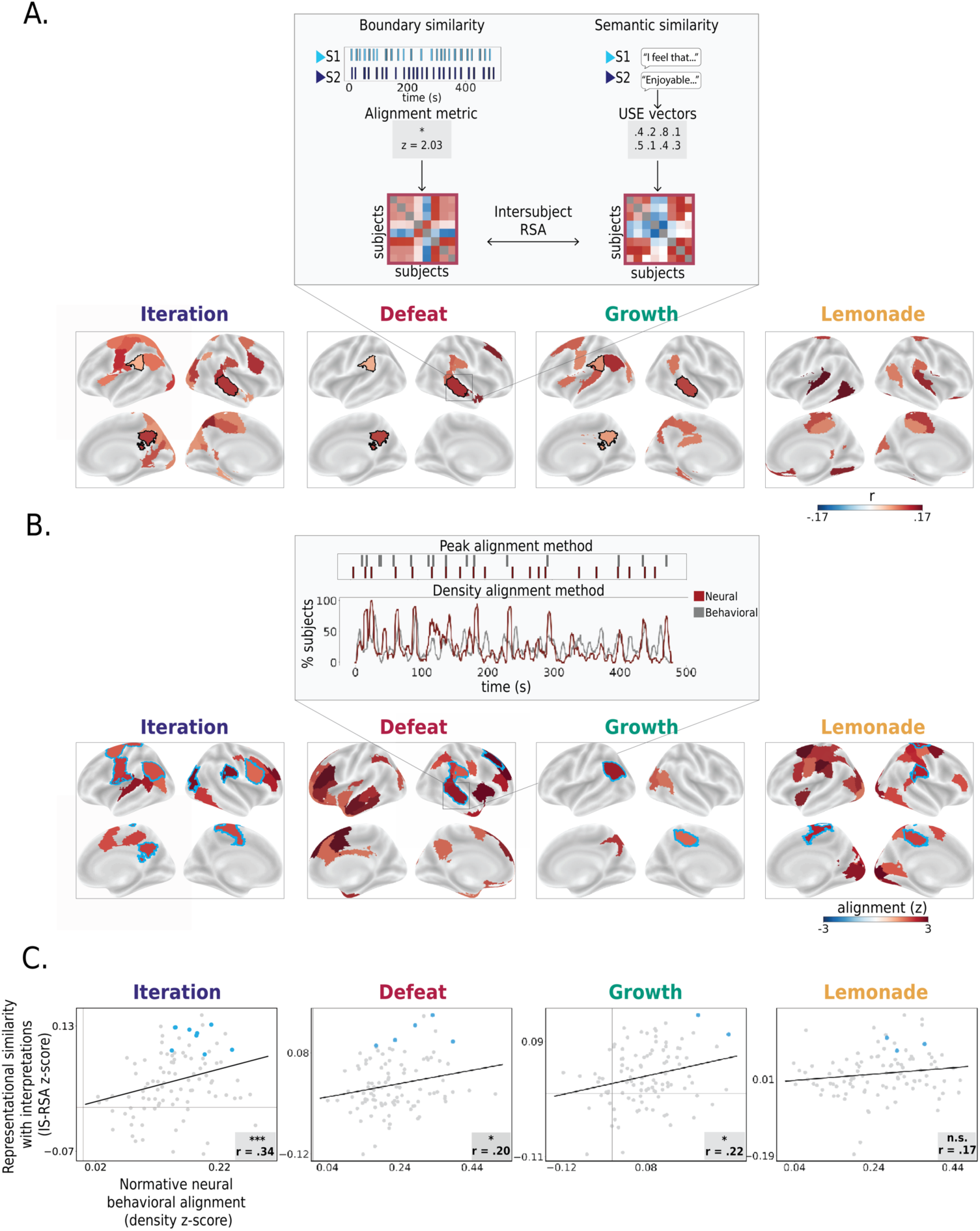
Relationship between neural event segmentation and behavior. **A. Inter-subject representational similarity between event boundary locations and ultimate appraisals.** Maps depict regions where pairs of subjects that were more similar in their neural event boundaries also had more similar interpretations of the movie (partial Mantel test controlling for objective memory performance; *q*<.05, FDR corrected to 100 regions within movie). Black contours indicate regions that showed significant representational similarity across all three social movies. Among these was a subregion of the superior temporal lobe (partial Mantel test, *q*<.05, corrected within movie), the precuneus, and the angular gyrus (partial Mantel test, *p*<.05 uncorrected across all three social movies). **B. Brain regions where normative neural boundaries align with normative behavioral boundaries.** Regions showing significant alignment between normative (group-level) behavioral and neural boundaries (*q*<.05, FDR corrected across 100 regions within movie). Importantly, though there are numerous regions where this relationship holds, specific regions vary across movies, suggesting that this relationship is somewhat movie-specific. Alignment between normative behavioral and neural boundaries was computed using two approaches (see *Methods*) (depicted for an example region in the superior temporal sulcus.) Blue contours indicate regions that also show significant alignment with appraisal (cf. Fig. 3A). **C. At the movie level, similar brain regions show relationships with both normative behavioral segmentation and individual-level appraisals.** The degree of alignment between neural and behavioral normative boundaries (“Density Alignment”) is correlated with the relationship between individual neural event boundaries and individual interpretations. Gray dots represent each region. Regions that show a significant relationship in both analyses (blue contours in 3B) are demarcated with blue dots. Values reflect Fisher z-transformed correlation coefficients. A Spearman correlation was used. Legend: * indicates *p*<.05;*** indicates *p*< .001.

Results (**Fig. 3A**) showed that in several brain regions, the degree of alignment in neural event boundaries during movie-watching predicted similarity in appraisal; i.e. subjects that were similar in event boundaries in these regions also tended to speak similarly about the movie afterwards. The specific regions showing these effects differed for each movie, but roughly corresponded to regions sensitive to each movie’s respective “high-level” content (ex: humans versus objects; *Iteration, Growth, Defeat* versus *Lemonade*; Speer et al. (2009). In the three movies with social information (*Iteration, Defeat* and *Growth*), these regions included high-order social cognition areas that are often associated with complex narrative processing, some of which are considered part of the default-mode network (DMN; for Review: Yeshurun et al. (2021)). Three regions— the superior temporal lobe, the angular gyrus and the precuneus—emerged in all three social movies (**Fig. 3A**; black contours). Significant regions in the non-social movie (*Lemonade*) had a different spatial pattern from the social movies. These regions included mostly somatomotor, dorsal-attention, and limbic regions, including a posterior temporal, ventral stream object-category selective region, the lingual gyrus, the temporal parietal junction (TPJ), cingulate, the orbitofrontal cortex, and a left somatomotor region previously linked to motor imagery (Chen et al. 2009; Chinier et al. 2014).

### Relationships between neural boundaries and interpretations are strongest in regions reflecting normative behavioral segmentation

As noted above, the different spatial patterns of relationships between neural event segmentation and appraisal across movies that we identified in the previous analysis are likely due to differences in movie content. For a particular movie, are the regions most closely tied to group-level behavioral segmentation during encoding also those most likely to show individual-difference relationships between segmentation and ultimate appraisal?

To answer this question, we must independently quantify the degree to which neural event boundaries in a region reflect behavioral event boundaries. While past work has shown that automatically detected neural event boundaries tend to align with behaviorally reported event boundaries in certain regions of association cortex such as the angular gyrus or the posterior medial cortex (Baldassano et al. 2017; Lee et al. 2021; Williams et al. 2022; Yates et al. 2022) this work has typically been limited to single stimuli, so the extent to which the regions showing behavioral-neural event alignment differ holds across stimuli has yet to be explored.

Towards this goal, we, again, used our auxiliary behavioral study (*Experiment 2*) and identified “normative” behavioral boundaries where subjects agreed, on average, that there was an event boundary. We then compared these normative behavioral boundaries to normative neural boundaries determined by fitting an HMM to the fMRI data averaged across subjects. We did this neural-behavioral normative boundary comparison using two approaches (**Fig. 3B** inset, see *Methods* section “Comparison of neural event boundaries and behavioral event boundaries”) and only considered regions that were significantly aligned to behavior (*p*<0.05) in both methods.

As expected, regions with significant neural-behavioral segmentation alignment varied across movies (**Fig. 3B**), but generally showed anatomical consistency with previous reports of regions that are sensitive to event changes (Zacks et al. 2001; Speer et al. 2003, 2007; DuBrow and Davachi 2016; Baldassano et al. 2017; Masís-Obando et al. 2022); namely, the temporoparietal junction (TPJ), prefrontal cortex (PFC), insula, parietal cortex, and cingulate (though note, interestingly, that *Lemonade*, the non-social movie, showed neural-behavioral segmentation alignment in earlier visual regions).

Finally, we correlated, across regions, the degree of cross-modal alignment between normative neural and behavioral boundaries (density metric; **Fig. 3B**) and the representational similarity r-value between individual neural event boundaries and appraisal (cf. **Fig. 3A**). Indeed, these two properties were positively correlated in three of the four movies (**Fig. 3C**), indicating that, for a given movie, segmentation in regions that mirror behaviorally reported event boundaries at the group level are also relevant for how that movie is ultimately remembered and appraised at the individual level. Regions that showed significant cross-modal normative alignment, as well as a significant relationship between individual boundary locations and appraisal, are highlighted with blue contours and blue dots in **Fig. 3B** and **3C**, respectively. We suggest that the same regions that show neural segmentation at moments corresponding to behaviorally reported boundaries were likely involved in appraising and translating these events into memory. Future work should investigate this in a more causal manner.

## Discussion

Individuals segment incoming information into events in different ways. Here, using four different short-film stimuli, we investigated individual differences in neural event boundaries and how they relate to behavior. Results showed that there is a posterior-to-anterior gradient for between-subject alignment in neural event boundaries that is tightly correlated with the rate of segmentation, such that regions that segment more slowly also show less alignment (i.e, more individual variability). Notably, we found strong movie effects for regions in the middle of this gradient: while alignment was high in low-level sensory regions and low in high-order narrative processing regions across movies, regions that are more tuned to a certain type of input showed variable alignment depending on the movie. We also describe one mechanism by which narratives may generate variable interpretations across people: individuals with more similar neural event boundaries in certain regions during movie-watching tended to have more similar appraisals of the movie.

Our first goal was to validate the use of automated event segmentation algorithms on fMRI data from individual subjects, and to characterize how much individuals vary in their neural event boundary locations. We found that although subjects show above-chance alignment in event boundaries in the majority of the cortex irrespective of the stimulus, the degree of alignment decreases from unimodal to transmodal association regions. This finding is in line with reports using inter-subject correlation (ISC) to show that synchrony of activity decreases from posterior to anterior regions (Yeshurun, Nguyen, et al. 2017; Haxby et al. 2020; Chang et al. 2022). Thus, regardless of measurement modality—ISC (continuous activity timecourses) or event segmentation (discrete state switches)— dynamic neural responses to external stimuli are more idiosyncratic in higher-order regions including the TPJ, STS, precuneus, dorsal medial and ventral medial prefrontal cortex, and the medial frontal gyrus. Notably, these regions are often considered part of the default mode network with reported involvement in social cognition, self-referential processing, and the consolidation of autobiographical memory (for review: (Yeshurun et al. 2021)).

We also found that the degree of alignment is tightly correlated with segmentation rate across regions. This broadens recent findings on event rate (Baldassano et al. 2017; Geerligs et al. 2022; Yates et al. 2022) and on the intrinsic process memory and temporal receptive windows (the length of time before a brain area’s response in which information will impact that response), which are all thought to follow a hierarchical posterior-to-anterior gradient as well (Lerner et al. 2011; Honey et al. 2012; Hasson et al. 2015). We suggest that information integration (as measured by event boundary alignment) is more stereotyped across subjects in sensory, unimodal cortex and becomes less standardized in higher-order regions with slower dynamics due, in part, to idiosyncratic processing strategies, experiences, and memories.

To support the link between idiosyncratic neural event boundaries and variable information integration, we demonstrate that patterns of neural event segmentation in certain regions relate to an individual’s memory of the segmented experience. Using ISC, it has been shown that individuals with shared context (Yeshurun, Swanson, et al. 2017), shared traits (e.g., paranoia; (Finn et al. 2018)), or shared experimentally manipulated perspectives (Lahnakoski et al. 2014) have similar neural responses while experiencing a narrative. We aimed to extend this past work and capture meaningful differences in how idiosyncratic neural activity during encoding relates to endogenously generated interpretations and recalls. Past work has theorized that current perceptual information interacts with long-term knowledge about event categories (schemas and scripts) when forming and updating event models (Radvansky and Zacks 2014). These stereotyped changes likely form shared boundaries (Baldassano et al. 2018; Masís-Obando et al. 2021) and reflect central hubs in the narrative (Lee and Chen 2022). We suggest that individual-specific boundaries—i.e., those not shared among the majority of subjects—may reflect moments with more idiosyncratic meanings (i.e., a moment activating one’s own autobiographical memory) that lead to variable interpretations of a stimulus. To test this, we leveraged complex narratives that generated variable appraisals across people and had subjects freely discuss the movies. We then used inter-subject representational similarity analysis to show that pairs of subjects with more similar event boundaries in numerous higher-order regions, including the angular gyrus, superior temporal lobe, and precuneus, also had more similar appraisals of each stimulus. A large subset of these regions also reflect behaviorally reported event boundaries. Altogether, these findings uphold the functional role of these regions in the high-level encoding of an experience and its ultimate behavioral consequences, including its organization into memory.

While the idea that stimulus features systematically affect group-level segmentation is not novel— (Newtson et al. 1977) were among the first to consider how a movie’s features encourage segmentation of continuous streams of information into units—here, we extend this to characterize how movie content affects neural and behavioral segmentation at the individual level. We demonstrated that although the spatial patterns of relative alignment were consistent across movies, the absolute degree of alignment differed with the specific movie and its content. While these four movies differed along numerous dimensions that could have affected degree of alignment, our two *a priori* hypothesized features—namely, the presence of both continuity editing and a social, character-driven, emotional narrative—drove both higher overall alignment across the cortex (cf. Fig. 2) and in behaviorally reported boundaries (cf. Fig. 3). There are some limitations to this work. First, although our choice of movies was somewhat principled in that it was based in existent theories about the effects of content on segmentation (Grall and Finn 2022), our ultimate stimulus set is still arbitrary and there are plenty of other low-, mid-, and high-level features that vary across these movies. We hope that through the release of our dataset, others can build on this work and investigate additional stimulus features that may affect individual segmentation patterns and variability therein. Second, our sample size (n=43) is relatively low for individual-differences work. However, a strength of our design is that we compared the same subjects across four movies. Furthermore, we did not attempt to link individual neural event segmentation patterns to trait-level behavioral measurements, an analysis for which we would likely be underpowered; rather, we focused on quantifying the overall degree of individual variability and how this differs as a function of both brain region and stimulus content, for which sample size should be a less limiting factor.

In sum, our work shows that individual differences in the location of neural event boundaries 1) meaningfully relate to behavior, and 2) vary with stimulus content in both their strength and spatial patterns, thereby emphasizing the importance of considering stimulus content in naturalistic neuroimaging paradigms. Future work should look at these relationships in terms of individual variability in the rate of segmentation. Furthermore, although numerous studies have explored individual differences in event segmentation at the behavioral level, here, we extend existing methods to identify individual differences at the neural level during encoding and their consequences for how a stimulus is ultimately remembered and appraised. Future work should further explore this relationship by identifying whether clinical or personality traits (see Zacks & Sargent (2010)) that are stable over time are associated with particular neural styles of segmentation that impact ongoing cognition and behavior.

## Acknowledgements

The authors thank Peter Bandettini, Peter Molfese, Daniel Handwerker, and Javier Gonzalez-Castillo for helpful discussions about experimental design and support with data collection. We thank Gang Chen for statistical advice and help implementing the LME-CRE analyses. We also thank Sofia Yawand-Wossen and Payton Weiner for their assistance with the movie labeling and content analysis, and the Undergrad Research Assistantships at Dartmouth program for funding their support.

